# Force-dependent stimulation of RNA unwinding by SARS-CoV-2 nsp13 helicase

**DOI:** 10.1101/2020.07.31.231274

**Authors:** Keith J. Mickolajczyk, Patrick M.M. Shelton, Michael Grasso, Xiaocong Cao, Sara E. Warrington, Amol Aher, Shixin Liu, Tarun M. Kapoor

## Abstract

The superfamily-1 helicase non-structural protein 13 (nsp13) is required for SARS-CoV-2 replication, making it an important antiviral therapeutic target. The mechanism and regulation of nsp13 has not been explored at the single-molecule level. Specifically, force-dependent unwinding experiments have yet to be performed for any coronavirus helicase. Here, using optical tweezers, we find that nsp13 unwinding frequency, processivity, and velocity increase substantially when a destabilizing force is applied to the dsRNA, suggesting a passive unwinding mechanism. These results, along with bulk assays, depict nsp13 as an intrinsically weak helicase that can be potently activated by picoNewton forces. Such force-dependent behavior contrasts the known behavior of other viral monomeric helicases, drawing stronger parallels to ring-shaped helicases. Our findings suggest that mechanoregulation, which may be provided by a directly bound RNA-dependent RNA polymerase, enables on-demand helicase activity on the relevant polynucleotide substrate during viral replication.

## MAIN TEXT

SARS-CoV-2 is a positive-stranded RNA coronavirus responsible for the current coronavirus 2019 (COVID-19) pandemic^1,2^. Its genome encodes sixteen non-structural proteins (nsp), including the RNA-dependent RNA polymerase nsp12 (RdRP), and the RNA helicase nsp13^3^. Nsp13 is essential for viral replication^4^ and was recently shown to form a complex together with nsp12^5^. Nsp13 from SARS-CoV-2 is a superfamily 1 helicase and is >99% similar to nsp13 from SARS-CoV-1^3^, differing by a single amino acid (V570I). This high level of conservation indicates that nsp13 from SARS-CoV-1 and -CoV-2 have a shared unwinding mechanism. It also indicates that antiviral therapeutics targeting the helicase would be effective towards SARS-CoV-1 and -CoV-2 nsp13, and potentially on helicases from future coronaviridae. Despite this therapeutic potential, the detailed mechanism and regulation of replicative helicases in coronaviridae in general remain poorly studied.

Previous work on SARS-CoV-1 nsp13 has established via bulk assays a 5’-3’ directionality of unwinding activity^6^. Nsp13 has been shown to unwind both RNA and DNA substrates with approximately equal initial rates, despite the virus having an RNA genome^7,8^. The efficiency of nsp13 unwindase activity is low compared to other helicases, but may be enhanced by either cooperativity or by interaction with the viral RdRp^8–10^. Consistently, recent structural data has shown that SARS-CoV-2 nsp13 can form a complex with RdRP^5^. Nonetheless, the potential mechanisms underlying nsp13 activation have not been explored.

Mechanical forces offer potential means for the regulation of viral helicase activation. Some viral helicases, such as hexameric T4 bacteriophage gp41 and T7 bacteriophage gp4, exhibit enhanced unwinding velocities when destabilizing forces are applied to their polynucleotide substrates^11,12^. Other viral helicases, such as the monomeric hepatitis C virus (HCV) nonstructural protein 3 (NS3) show little change in unwinding velocity when forces are applied to their substrate^13^. Strong force-dependence of velocity is correlated with a passive unwinding mechanism (i.e. energy from ATP hydrolysis is used for translocation, but not for breaking base pairs), whereas weak force-dependence of velocity is correlated with an active unwinding mechanism (i.e. energy from ATP hydrolysis is used to directly separate base pairs)^14–16^. As such, force-dependence of the unwinding activity of a helicase has fundamental implications for its regulation during physiological operation. Force-dependence measurements have yet to be performed for any coronavirus helicase.

In the current work, we investigate the mechanism of SAS-CoV-2 nsp13-driven polynucleotide unwinding activity using both bulk and single-molecule approaches. Using bulk assays, we find that nsp13 unwinds both DNA and RNA with weak intrinsic activity. Using optical tweezers, we find that applying picoNewton forces, as could be provided by a bound RdRp to the RNA substrate increases the event frequency, unwinding velocity, and processivity of nsp13, leading to a >50-fold enhancement of catalytic activity. These results indicate that nsp13 utilizes a passive unwinding mechanism, and identifies mechanoregulation as a sufficient means of nsp13 activation.

## RESULTS

### Biochemical characterization of SARS-CoV-2 nsp13

The SARS-CoV-2 nsp13 helicase consists of an N-terminal zinc binding domain (ZBD), followed by stalk and beta domains (S and 1B domains, respectively), and finally the catalytic RecA1 and RecA2 (A1 and A2) helicase domains (**Fig. 1a**). Previous structural and biochemical data on SARS-CoV-1 and MERS-CoV^4,17^ have identified the ZBD and S domains as important for structural stability of the enzyme. The A1 and A2 domains regulate ATP hydrolysis, and, along with the B1 domain, are proposed to regulate oligonucleotide binding.

**Figure 1.**
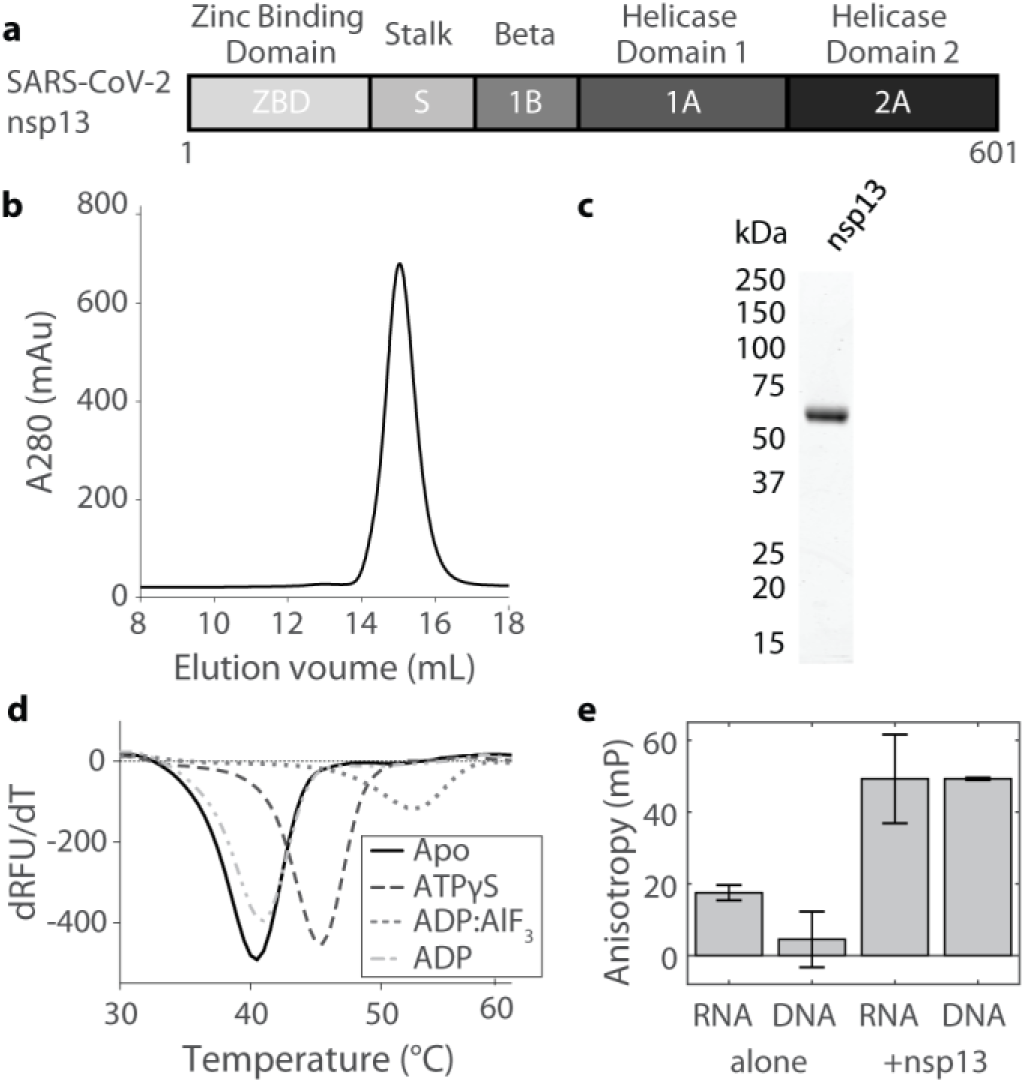
Biochemical characterization of nsp13. a. Diagram showing the domain architecture of SARS-CoV-2 nsp13. b. Size exclusion chromatography (Superdex 200 Increase) of purified nsp13. c. SDS-PAGE gel (Coomassie stain) showing nsp13 purity. d. Differential scanning fluorimetry (DSF) of SARS-CoV-2 nsp13 in the absence and presence of ATP analogs (1 mM). Melting temperatures are: APO- 40.5 °C, ATPγS- 45.5 °C, ADP:AlF_3_- 52.5 °C, and ADP- 41.0 °C for n=2 independent measurements from two separate protein preparations. e. Fluorescence anisotropy measurements for fluorescence DNA and RNA partial duplexes (10 nM) with and without nsp13 (3.4 μM) added. Data shown as mean standard ± deviation (SD) for n=2 independent measurements from two separate protein preparations.

Biochemical studies of SARS-CoV-1 nsp13 indicated that a GST-tag, which has the potential to induce dimerization, compared to a His_6_-tag, could substantially alter activity (>280-fold)8. However, the positively-charged His_6_-tag may also artificially bias polynucleotide binding. Therefore, for our analyses we used recombinant affinity tag-free, full-length SARS-CoV-2 nsp13 (hereafter, nsp13), which we have reported recently^5^. Briefly, the protein was subjected to Ni-NTA affinity, cation exchange (CaptoS HP) and size exclusion (S200 increase) chromatography. The addition of an ion exchange step reduced DNA contamination and a monodisperse peak off of the gel filtration column at 15 mL (**Fig. 1b**), and SDS-PAGE (Coomassie) analysis revealed >95% purity (**Fig. 1c**).

To examine the nucleotide binding capabilities of nsp13, we used differential scanning fluorimetry (DSF) to measure shifts in melting temperature^18^. We found that nsp13 alone (apo) had a melting temperature of 40.5 °C, and that the addition of ATPγS or ADP:AlF_3_ (1 mM) shifted the melting temperature ~5-12°C (**Fig. 1d**). These shifts indicate stabilization via ligand binding, and are consistent with recent structural data showing ADP:AlF_3_ binding within the nsp13 active site^5^ (**Fig. 1d**). No significant shift was observed in the presence of ADP (1 mM), indicating minimal protein stabilization (**Fig.1d**).

To examine nsp13 binding to RNA and DNA, we performed fluorescence anisotropy measurements. We designed fluorescently-tagged double-stranded DNA and RNA substrates containing an 18-bp duplex region with a 10-nucleotide 5’ overhang. A 10-nucleotide overhang allows only one copy of nsp13 to bind at a time^9^. The fluorescence anisotropy of these substrates (10 nM) was measured both in the absence and presence of nsp13 (3.4 μM) in nucleotide-free buffer. We measured an increase in fluorescence anisotropy for both DNA and RNA, indicating that nsp13 can bind both of these substrates (**Fig. 1e**). Together, these data indicate that recombinant tag-free nsp13 binds nucleotide analogs, and can bind both RNA and DNA in the absence of nucleotide.

### Unwinding activity of nsp13 on DNA and RNA substrates

To examine the helicase activity of SARS-CoV-2 nsp13, we utilized a bulk fluorescence-based single-turnover assay^19^. Briefly, we synthesized the DNA and RNA substrates described above but with the addition of a quencher on the previously unlabelled strand such that fluorescence signal is suppressed while in the duplex, but not when the strands are dissociated (**Fig. 2a**). In this assay, nsp13 (10 nM) unwound DNA in the presence of MgATP (2 mM), as revealed by an increase in the fluorescence signal (**Fig. 2b**). This activity was also observed using a gel-based assay (**Fig. S1**). We found that helicase activity could be readily observed at nsp13 concentrations below 100 pM in the presence of DNA (1 μM) (**Fig. 2c**). We next performed an initial velocity analysis of nsp13 and observed unwinding rates that increased with DNA concentration. Fitting these data to the Michaelis-Menten equation revealed a *K*_M_ of 1.05 ± 0.15 μM for DNA and a *k*_cat_ of 0.53 ± 0.03 s^−1^ (fit ± 95% confidence interval; **Fig. 2d**). This rate of DNA unwinding is similar to that reported for HCV NS3 helicase in a fluorescence-based helicase assay^20^. We determined a Hill coefficient of ~1, indicating a lack of cooperativity in nsp13-driven unwinding for our substrates.

**Figure 2.**
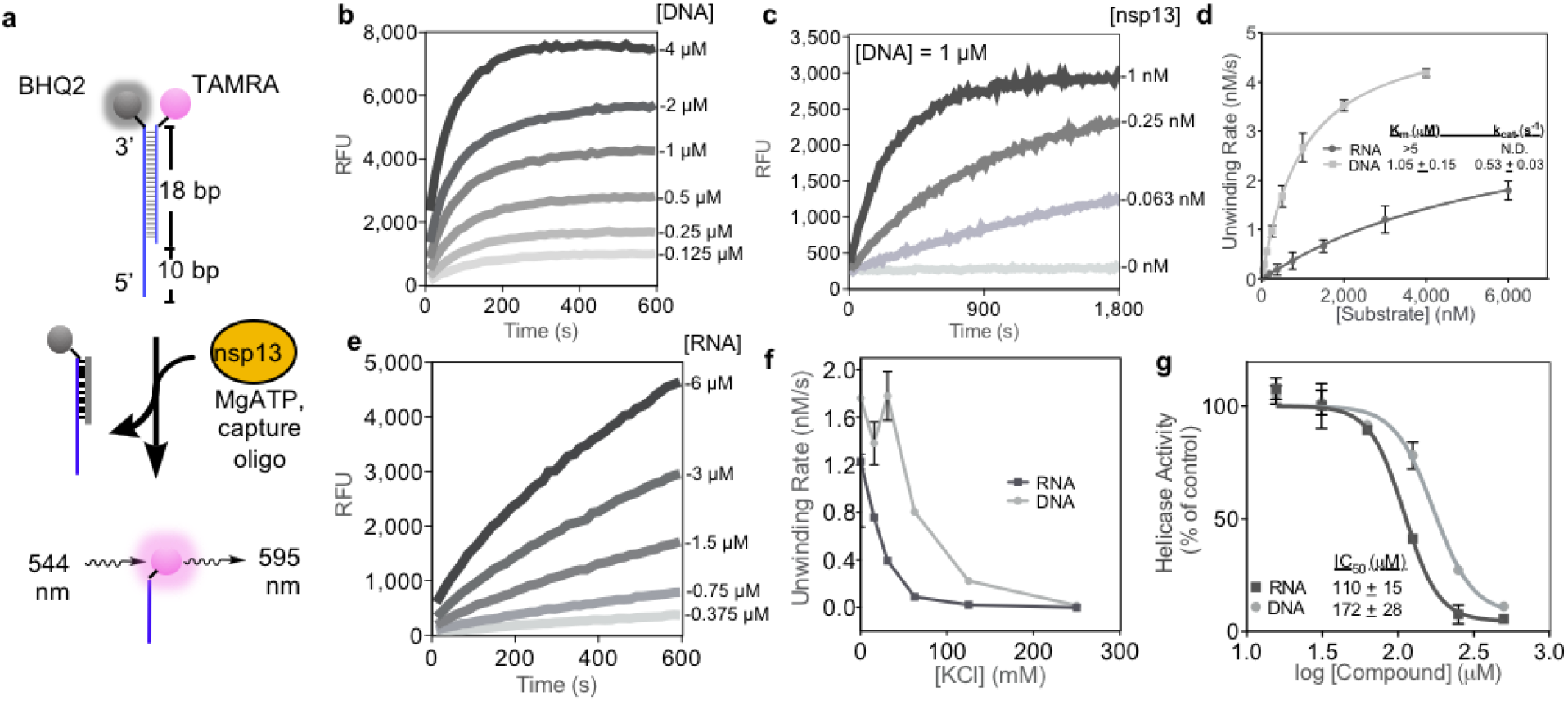
Nsp13 is a DNA and RNA unwindase. a. Diagram of fluorescence-based helicase assay using a partial duplex oligonucleotide substrate with 10-nt overhang. Unwinding of the duplex results in a fluorescent signal. b. Nsp13 helicase activity (10 nM) as a function of time with increasing DNA substrate concentration. c. Nsp13 helicase activity as a function of time with a constant concentration of DNA (1 μM) and increasing nsp13 concentrations. d. Plot of initial substrate unwinding velocities versus substrate concentration (0-4 μM for DNA, 0-6 μM for RNA). Data points show the mean ± SD for n = 2 or 3 measurements. Fitting the data points to the Michaelis-Menten model revealed K_M_ and k_cat_ values for DNA unwinding (inset; fit ±95% confidence intervals). e. Nsp13 helicase activity (10 nM) as a function of time with increasing RNA substrate concentration. f. Initial DNA unwinding rates at varying KCl concentration for DNA and RNA substrates. Data points show the mean ± SD for n = 2 measurements. g. Dose-response curves of nsp13 helicase activity in the presence of increasing concentration of ADP:AlF_3_ (0-500 μM). Data points show the mean ± SD for n = 3 measurements. IC_50_ values were determined by fitting the data to the Hill equation (inset).

We next examined nsp13 helicase activity using an RNA duplex with an analogous sequence. As with DNA, we observed a substrate concentration-dependent increase in fluorescence signal (**Fig. 2e**). Compared to DNA, we observe that the unwinding rate for the RNA substrate is ~2-3-fold lower, and that the initial rate profile for RNA remains linear even at the highest substrate concentration (6 μM) tested (**Fig. 2d, e**). Therefore, we can only estimate a *K*_M_ > 5 μM for RNA.

To examine the basis of the difference between DNA and RNA unwinding in the helicase assay, we considered that ionic strength (i.e. KCl concentration) could alter nsp13 duplex binding affinity due to the electrostatic nature of enzyme-nucleic acid interactions. DNA unwinding was faster than RNA (between 1.5 to 8-fold) at all salt concentrations tested (**Fig. 2f**). Nsp13 demonstrated the highest activity when no KCl was added to the assay, and increasing the salt concentration led to a decrease in duplex unwinding rate, with no measurable activity at ≥250 mM KCl. A similar salt dependence of nsp13 helicase activity was observed in gel-based assays (**Fig. S2**). As a control, ADP:AlF_3_ inhibited both nsp13-dependent RNA and DNA (500 nM) unwinding with similar IC_50_ values (**Fig. 2g**). Overall, our data suggest that nsp13 possesses 5’-3’ helicase activity, albeit with a weak K_M_, and, for our specific substrates, may unwind DNA more readily than RNA.

### Single-molecule measurements of nsp13-driven RNA unwinding

To further investigate the RNA helicase activity of nsp13, we designed an optical trap assay to directly visualize RNA unwinding at the single-molecule level. Briefly an RNA hairpin including a 20-nucleotide 5’ loading zone, a 180-bp duplex region, and a tetraloop, was annealed to two 1.5-kbp dsDNA handles, which in turn were conjugated to polystyrene beads (see Methods; **Fig. 3a**). One bead was held steady in a micropipette, while the other was held in an optical trap and manipulated to apply a specified amount of force. A single tether between the two beads was ensured by measuring the force-extension curve of each tether (**Fig. 3b**). Under 20 pN, the hairpin remained stably folded by itself for the duration of the experiments. Nsp13 was then injected into the sample chamber, and dsRNA unwinding was monitored by the necessary changes in the optical trap position required to hold a constant force. Nsp13-catalyzed unwinding of the hairpin would result in an increase in the distance between the two beads.

**Figure 3.**
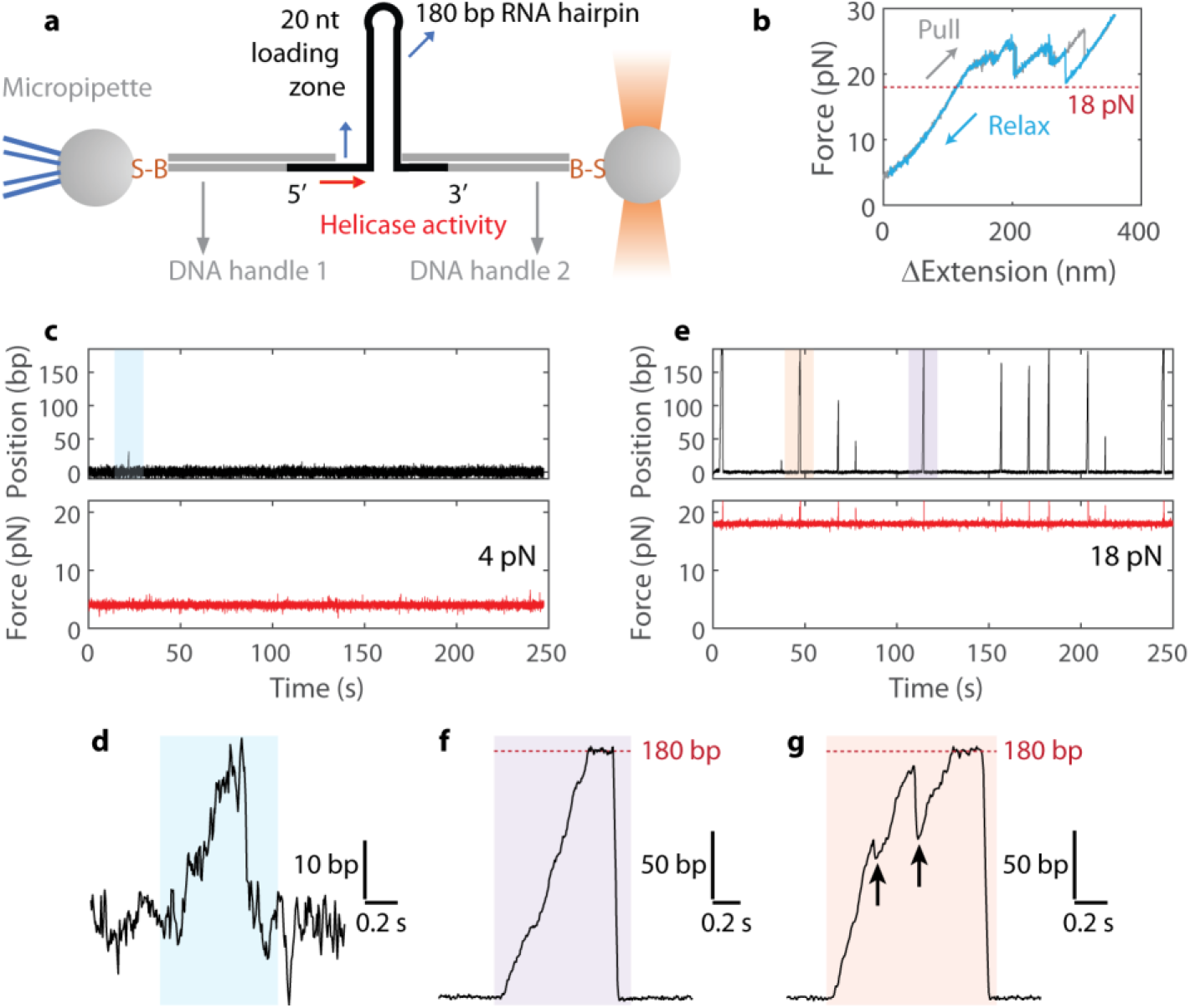
Single-molecule measurement of nsp13 helicase activity. a. Assay geometry showing dsRNA hairpin complex. dsDNA handles separate the hairpin from the beads (connected by biotin-streptavidin, B-S). A 20 nucleotide (nt) region allows single nsp13 molecules to bind and begin 5’-3’ helicase activity. Motion of the optically-trapped bead (top) under force-feedback enables a set force to be applied to the dsRNA hairpin substrate. Diagram not to scale. b. Representative force-extension curve of a single DNA/RNA tether (no nsp13 present). Above 20 pN, external force begins to unwind the RNA hairpin. c. Representative example trace at 4 pN. Only one event is observed in ~250 s. d. Zoomed-in blue region from panel c. e. Representative example trace at 18 pN. Numerous events are observed in ~250 s. f. Zoomed-in orange region from panel e. Arrows denote slippage events. Red dotted line denotes 180 bp, the total length of the RNA hairpin. g. Zoomed-in purple region from panel d. Red dotted line denotes 180 bp, the total length of the RNA hairpin.

We first made measurements at low applied forces (4 pN). At this force, the unwinding activity of nsp13 was consistent with the low catalytic efficiency measured in the bulk assays (**Fig. 2**). The long time period between the unwinding events relative to the durations of these events suggest that each unwinding is likely driven by a single helicase enzyme^11^. RNA unwinding events by nsp13 were rare and short, typically terminating after a few tens of bp, indicating poor processivity (**Fig. 3c, d**). We next made measurements at 18 pN, the highest applied force feasible without starting to open the hairpin (**Fig. 3b**). We observed three major differences at 18 pN (**Fig 2e**) in comparison to 4 pN: (i) unwinding events occurred more frequently at the same nsp13 concentration, (ii) unwinding events became longer in distance, and (iii) unwinding appeared to be much faster. In some instances, nsp13 unwound the entire 180 bp hairpin (**Fig. 3f**). We also observed slippage events in ~33% of recorded traces at 18 pN; These events were marked by sudden (>1 kbp-s^−1^) backward motions along the RNA followed by a rapid resumption in unwinding (**Fig. 3g**). Furthermore, the unwinding velocity and processivity decreased with reduced ATP (**Fig. S3**), indicating that nsp13 may detach from the RNA while waiting for ATP.

### Nsp13 event frequency, processivity, and velocity are enhanced by applied force

To quantify the force-dependence of nsp13 helicase activity, we made single-molecule measurements of dsRNA unwinding at multiple applied forces between 4 and 18 pN (**Fig. 4a**). We first measured the event frequency, or number of independent unwinding events per unit time. The event frequency increased ~14-fold between 4 and 18 pN (**Fig. 4b**). Assuming a simple model where there is no rate-limiting step to initiate unwinding, the event frequency is proportional to the on-rate of nsp13 binding to RNA, meaning that applied force on the RNA substrate may increase the nsp13 on-rate ~14-fold. The low event frequency extrapolated to zero force is consistent with the weak *K*_M_ measured in bulk assays (**Fig. 2d**).

**Figure 4.**
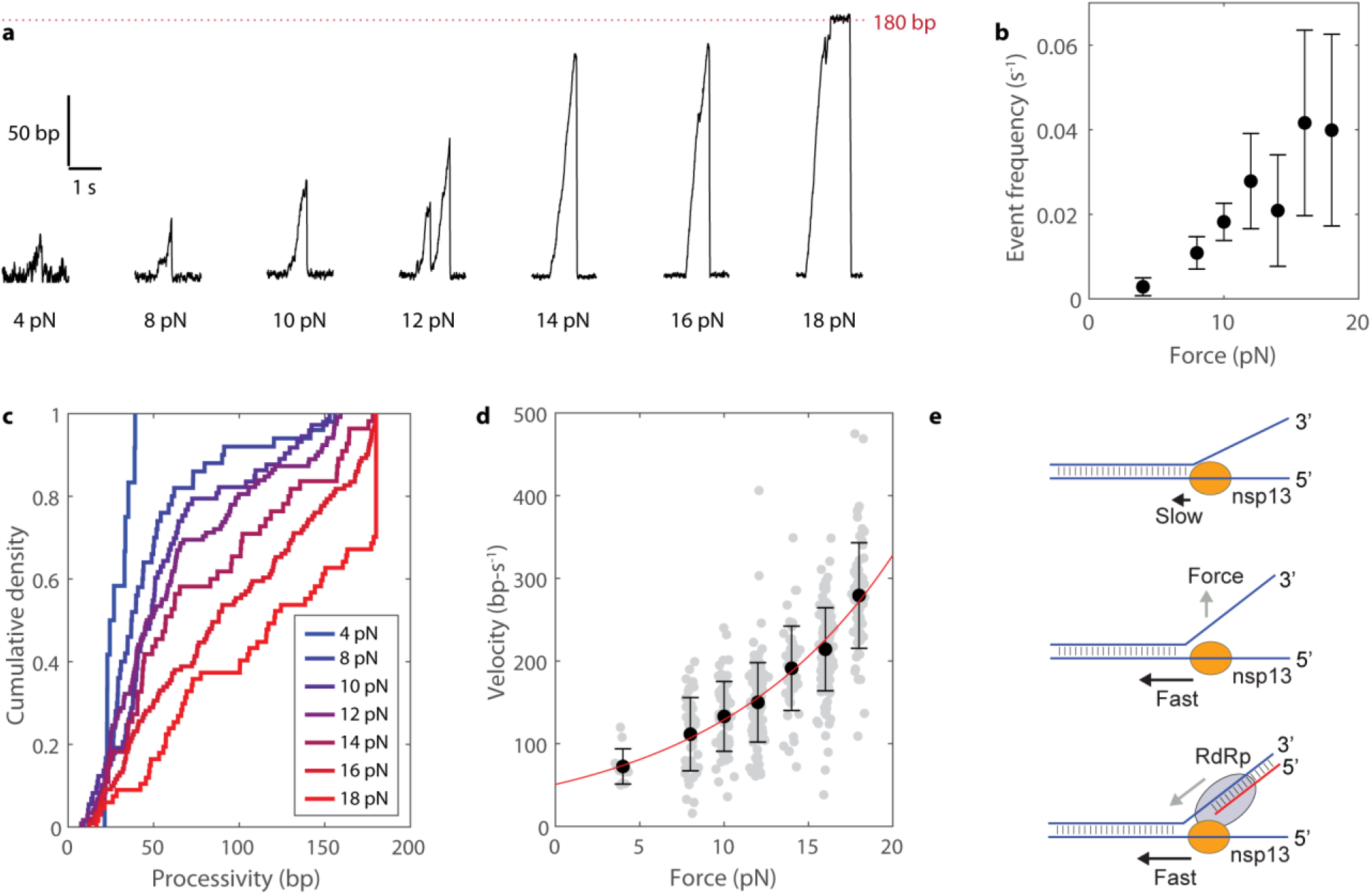
Force-dependent activation of nsp13 dsRNA helicase activity. a. Example traces of nsp13 unwinding dsRNA at 4, 8, 10, 12, 14, 16, and 18 pN force. b. The number of new processive events per second at various forces. Data shown as mean ± SD for n=5-10 independent RNA hairpins. c. Distributions of measured processivities for nsp13 at various forces (n=12, 50, 73, 118, 55, 121, 67 events for 4, 8, 10, 12, 14, 16, and 18 pN applied force, respectively). 180 bp is the total length of the RNA hairpin. d. The measured velocity of nsp13 at various forces (n=12, 50, 73, 118, 55, 121, 67 events for 4, 8, 10, 12, 14, 16, and 18 pN applied force, respectively). Each measurement shown as a gray dot slightly spaced out in force for visibility. Black data points show mean ± SD. Data were fitted to an exponential function *v*(*F*) = *v*_0_*exp*(*F*/*F_e_*) (red line), revealing a Y-intercept (v_0_) of 51 ± 9 bp and a rate term (F_e_) of 10.4 ± 1.3 pN (fitted values ± 95% confidence intervals). e. Model showing that external force on the dsRNA substrate enhances the velocity of nsp13. One potential source of force is the RNA-dependent RNA polymerase.

We next measured the processivity, or total length of dsRNA unwound during each event. The processivity increased monotonically from 4 pN, where no event was greater than 50 bp, to 18 pN, where ~35% of the nsp13 molecules unwound the entire 180 bp dsRNA duplex region (**Fig. 4c**). At the lower forces measured (4-14 pN), where events were not truncated by the limited length of the RNA hairpin, distributions of measured processivities could be fitted to a single exponential, indicating that, for this given RNA hairpin, nsp13 has a constant probability of detaching as it unwinds (i.e. does not preferentially detach at a sequence-specific position) (**Fig. S4**).

We next measured the velocity of nsp13 in response to applied force on the dsRNA substrate. Remarkably, we found that the velocity increased exponentially with force, increasing from ~70 bp-s^−1^ at 4 pN to ~280 bp-s^−1^ at 18 pN (**Fig. 4d**). Velocity is a measure of *k*_cat_, meaning that applied force increases the *k*_cat_ ~4-fold. We can also estimate the nsp13 off-rate as the velocity divided by the processivity. As both of these values approximately increase in tandem over the measured force range, we estimate the nsp13 off-rate to be force-independent at ~2 s^−1^ (**Fig. 4c, d)**. Coupling the 4-fold change in the *k*_cat_ with the 14-fold change in the *K*_M_, we estimate that increasing the applied force for 4 to 18 pN increases the catalytic efficiency >50-fold. Fitting the velocity data to an exponential function yielded an extrapolated zero-force velocity of ~50 bp s^−1^ (**Fig. 4d**). Such scaling of the velocity indicates that nps13 likely follows a passive helicase mechanism^14,15^. Overall, we find that applying force to the dsRNA substrate enhances nsp13 unwinding frequency, processivity, and velocity.

## DISCUSSION

In the current work, we show that SARS-CoV2 nsp13 has weak intrinsic helicase activity that can be stimulated >50-fold by externally provided forces. These results provide insight into the mechanism and regulation of nsp13, enabling a deeper molecular understanding of the processes underlying SARS-Cov2 replication.

While the *in vivo* velocity of nsp13 has not yet been determined, the *E. coli* replication complex travels at ~1,000 bp s^−1 14,21^. We thus speculate that nsp13 moves much faster than its basal unwinding velocity during viral infection. Our data suggest that nsp13 RNA helicase activity can be stimulated >50-fold by forces in the 10-20 pN regime that act on the nucleic acid substrate **(Fig 4e)**. *E. coli* RNA polymerase and T7 viral RNA polymerase can each provide ~20 pN of force^22,23^. Therefore, we speculate that simultaneous action of the RdRp on the same duplex may provide the external force on the RNA substrate necessary for activated nsp13 unwinding (**Fig. 4e**). While the recent structural data does not directly inform on how nsp13 and the RdRp together work on the same RNA substrate^5^, existing kinetic data shows that RdRP enhances nsp13 helicases activity^8^. Moreover, there are numerous instances of polymerases activating helicases in other viral systems, including Zika^24^, T7^12,21,25^, and T4^26–28^. While posttranslational modifications or induced conformational changes may also contribute to nsp13 regulation, we show here that external force on the dsRNA substrate is sufficient to activate the helicase.

Replicative helicases from the six superfamilies^29^ fall into two broad classes: ring-shaped hexameric helicases and non-hexameric helicases^30,31^. Functionally, two mechanisms have been proposed for helicases to date: an *active* mechanism, in which the helicase directly destabilizes the DNA or RNA substrate, and a *passive* mechanism, in which the helicase opportunistically advances upon the spontaneous opening of base pairs^14–16^. These mechanisms are not mutually exclusive, and a continuum between them likely exists^32^. Nsp13 is non-hexameric^5^, and based on the shape of the force-velocity curve (**Fig. 4d**), we propose that it more closely matches the definition of a passive helicase. Notably, a passive mechanism has, to our knowledge, not been characterized for a non-hexameric helicase to date. The non-hexameric superfamily 2 helicase HCV NS3, and the non-hexameric superfamily 1 DNA repair helicase UvrD are proposed to be active helicases, each having unwinding velocities that do not increase in response to destabilizing forces^13,33^. We show here that the unwindase activity of nsp13 more closely matches that of ring-shaped bacteriophage DNA helicases such as T4 gp41 and T7 gp4, each of which show a strong force-dependent velocity^11,12^. The similarities between the velocities and processivities between nsp13 and T4 gp41 specifically, both basal and in response to external forces, suggest a common mechanism^11,34^.

Direct force-activation on nsp13 unwindase activity enables on-demand regulation. This is likely important to viral RNA genome stability, as promiscuous unwinding would expose it to degradation. A default low-activity state for the helicase also maintains the integrity of the viral replication machinery and prevents dissociation^35^. Nsp13 in SARS-CoV-2 is >99% similar by sequence identity to its counterpart in SARS-CoV-1^3^, suggesting that force-regulation of replicative helicase activity is likely conserved across coronaviridae that are evolutionarily proximal to the causative agent of COVID-19.

The high conservation of nsp13 helicases across coronaviridae suggests that therapeutic agents could be developed to treat COVID-19 and other diseases. Targeting viral helicases in general has been challenging^36,37^. However, recent successes in targeting different ATPases^38,39^ suggests that selective and potent inhibitors of nsp13 could be developed. The bulk and single-molecule assay platforms reported here may be leveraged in future drug discovery efforts and in detailed mechanistic investigation of nsp13 inhibition.

## MATERIALS AND METHODS

### Cloning

The nsp13 plasmid was generated as previously described ^5^. Briefly, the OptimumGene method (Genscript) was used to optimize the full-length SARS-CoV-2 nsp13 helicase sequence for insect cell expression. The gene was synthesized and cloned into the pet28(a)+ vector, using NdeI and XhoI restriction sites, by Genscript. We then performed mutagenesis using the Quikchange mutagenesis kit (Agilent) to insert an HRV-3C (PreScission) protease cleavage site and remove the thrombin cleavage site.

### Protein Expression and Purification

Nsp13 was expressed and purified as reported previously with minimal changes ^5^. Briefly, the plasmid expressing the SARS-CoV-2 nsp13 helicase (SARS-CoV-2-nsp13 Pet28a or SARS-CoV-2-nsp13 Pet28a-PreScission) was transformed into BL21 Rosetta cells and used to make a starter culture of 100 mL of LB with 100 μL of 50 mg/mL kanamycin. The starter culture was grown overnight at 37 °C. The next day, 4-6 L of LB were inoculated with 10 mL of SARS-CoV-2-nsp13 Pet28a-PreScission starter culture per L. These cultures were grown at 37 °C for ~2 hrs, or until an OD_600_ of 0.6. The temperature was then lowered to 18 °C and once at temperature, each L culture was induced with 1 mL of 0.2M IPTG per liter of growth. The cultures were left overnight (~16 hrs) at 18 °C and harvested the next day by spinning down at ~5000 g for 10 minutes and resuspend pellets in affinity buffer (50 mM Hepes pH 7.0, 500 mM NaCl, 4 mM MgCl_2_, 5% Glycerol, 1 mM PMSF, 20 mM Imidazole, 1 mM ATP, 1 mM BME). 9 μL of benzonase was added to the resuspended pellets and the pellet was subsequently lysed using an Avestin Emulsiflex C5 with pressure exceeding 10,000 psi. The lysate was then centrifuged using a Beckman Coulter ultracentrifuge and a Type 70 Ti rotor at 15,000 rpm for 45 minutes at 4 °C. The supernatant was then added to 2 mL of Ni-NTA resin equilibrated with affinity buffer and incubated at 4 °C with rocking for 1 hr. The supernatant and resin mixture was then passed through a gravity column, and the resin was subsequently washed with ~900 mL of affinity buffer. Once complete, the protein was eluted with 15-20 mL of affinity buffer + 250 mM imidazole. 100 uL of prescission protease (~4 mg/mL) were added to the elution and the protein was left to dialyze overnight in dialysis buffer (50 mM Hepes pH 7.0, 100 mM NaCl, 4 mM MgCl_2_, 1 mM TCEP). The elution is diluted 2Xfold with milliQ-H_2_O the next morning to give ion exchange A buffer (25 mM Hepes pH 7.0, 50 mM NaCl,2 mM MgCl_2_, 0.5 mM TCEP) and loaded onto a CaptoS HP column (Cytiva) at a flow rate of 1 mL/min and then eluted with a gradient from 0% ion exchange B buffer (25 mM Hepes pH 7.0, 1 M NaCl, 2 mM MgCl_2_, 0.5 mM TCEP to 50% ion exchange B buffer over 30 minutes. The protein eluted at around ~5-10% ion exchange B buffer. The protein was then concentrated to 500 μL using a 50 kDa cutoff Amicon Ultra-4 Centrifugal filter (Millipore UFC805008) and injected onto a Superdex 200 increase gel filtration column in sizing buffer (25 mM HEPES pH 7.0, 250 mM KCl, 1 mM MgCl_2_, 5% glycerol, 1 mM TCEP). The protein was flash frozen in liquid N_2_ and stored at −80 °C.

### Differential Scanning Fluorimetry

These experiments were carried out on a C1000 Touch Thermal cycler CFX-96 instrument (GE Healthcare). recombinant constructs were diluted to a final concentration of 4 μM in DSF buffer (25 mM HEPES-KOH pH 7.5, 25 mM KCl, 5 mM MgCl_2_, 1 mM TCEP, 0.005% triton-X) and assayed in a 96-well plate (Hard-shell HSP9665 Bio-Rad). Ligands were added to a final concentration of 1 mM. Sypro Orange fluorescent dye (Sigma S5692, excitation 490 nm, emission 590 nm) was used at 1:500 dilution. The temperature was linearly increased with a step of 0.5 °C for 55 minutes, from 25 °C to 95 °C and fluorescence readings were taken at each interval. Melting temperatures were recorded as the minimum value of the first derivative of the fluorescence vs temperature curves. These experiments were

### Preparation of DNA and RNA Duplex Substrates

All oligos used for the preparation of DNA and RNA substrates were purchased from Integrated DNA Technologies (IDT). The fluorescently-quenched partial duplex DNA substrate was prepared by mixing Oligo1 (5’-TTTTTTTTTTCTGATGTTAGCAGCTTCG-BHQ2-3’) and Oligo2 (5’-TAMRA-CGAAGCTGCTAACATCAG-3’) in a 1.2:1 molar ratio and annealed by heating at 95 °C for 5 min in a temperature controlled aluminum block, followed by cooling the sample at a rate of 1 °C/min until reaching 25 °C. The fluorescent partial duplex DNA substrate (lacking BHQ2) was prepared by mixing unlabeled Oligo1 and Oligo2 in a 1.2:1 molar ratio and annealing as above. The fluorescently-quenched partial duplex RNA substrate was prepared by mixing Oligo3 (5’-UUUUUUUUUUCUGAUGUUAGCAGCUUCG-BHQ2-3’) and Oligo4 (5’-TAMRA-CGAAGCUGCUAACAUCAG-3’) in a 1.2:1 molar ratio and annealed by heating at 75 °C for 5 min in a temperature controlled aluminum block, followed by cooling the sample at a rate of 1 °C/min until reaching 25 °C. The fluorescent partial duplex RNA substrate (lacking BHQ2) was prepared by mixing unlabeled Oligo3 and Oligo4 in a 1.2:1 molar ratio and annealing as per the fluorescently-quenched partial duplex RNA substrate. Capture oligos, complementary to the duplexed portion of the BHQ2 strands, used in helicase assays for DNA and RNA substrates, have the sequence 5’-CGAAGCTGCTAACATCAG-3’ and 5’-CGAAGCUGCUAACAUCAG-3’, respectively.

### Fluorescence Anisotropy Binding Assay

For fluorescence anisotropy studies, nsp13 (34 μM, in buffer containing 20 mM HEPES (PH=7.5), 5 mM MgCl_2_, 150 mM KCl, 1 mM TCEP, and 3% glycerol) and partial duplex DNA/RNA (as prepared above; 100 nM in 20 mM HEPES (PH=7.5), 5 mM MgCl_2_, 150 mM KCl, 1 mM TCEP) were diluted 10x into assay buffer (20 mM HEPES (PH=7.5), 5 mM MgCl_2_, 1 mM TCEP, 0.005% Triton X100) with KCl added to a final concentration of 25 mM (20 μL final reaction volume). Fluorescence anisotropy was measured after incubating for ten minutes at room temperature using a Synergy Neo2 plate reader (Biotek; 532/25 nm excitation filter, 590/35 nm emission filter) in dual photomultiplier tube (PMT) mode. Gain values on the two PMTs (reading parallel and perpendicular emission) were automatically scaled to give equal counts from a DNA or RNA-fluorophore-only control well, and the G factor was automatically adjusted.

### Microplate-based Helicase Activity Assays

Initial velocities of nsp13 helicase were measured by mixing recombinant nsp13 (10 nM) with partial duplex DNA or RNA substrates (0-6 μM) in the presence of ATP (2 mM) and capture oligo (8 or 12 μM) in 24 μL reaction volumes using solid black 384-well NBS microplates (Corning). Nsp13 was prepared as a 2x stock solution in helicase buffer (20 mM HEPES, pH 7.5, 40 mM KCl, 5 mM MgCl_2_, 0.5 mM EDTA, 1 mM TCEP, 0.5 mM glutathione, 0.01% triton X100 and 0.1 mg/mL BSA) and 12 μL was added to reaction wells. DNA/RNA duplex substrates were prepared as 4x stock solutions in helicase buffer followed by 2-fold serial dilutions in helicase buffer. Substrate dilutions (6 μL) were added to reaction wells containing nsp13 and incubated for 10 min. ATP/capture oligos were prepared as a combined 4x stock solution in helicase buffer, and 6 μL were added to reaction wells to initiate the reactions. Control reactions included wells with no enzyme and no ATP added. Substrate unwinding was measured by continuously recording the fluorescence (excitation: 544 nm, emission: 590 nm) on a Synergy Neo2 plate reader (Biotek) for 30 min, 15 s intervals. Relative fluorescence units (RFU) were converted to units of molarity with standard curves generated from DNA Oligo2 and RNA Oligo4. Enzyme parameters K_M_, k_cat_ and Hill coefficients (h) were determined by fitting the plot of initial rates (first ~150 s) against substrate concentration to the Michaelis-Menten equation or the Hill equation using Prism v.6.0 (GraphPad Software, Inc).

Variable enzyme concentration assays were performed by combining a 4x substrate/capture oligo (6 μL) stock solution in helicase buffer with 2-fold serial dilution of 2x nsp13 (12 μL) in helicase buffer in reaction wells. Reactions were initiated by the addition of a 4x ATP solution in helicase buffer. Final concentrations were 0-2 nM nsp13, 0.5 μM DNA/RNA substrate, 2.5 μM capture oligo and 2 mM ATP. Control reactions containing no ATP were included.

Variable KCl concentration assays were performed by combining a 4x nsp13/substrate/capture oligo (6 μL) stock solution in helicase buffer (-KCl) with 2-fold serial dilutions of 2x KCl solutions (12 μL) in helicase buffer in reaction wells. Reactions were initiated by the addition of a 4x ATP solution in helicase buffer. Final concentrations were 10 nM nsp13, 0.5 μM DNA/RNA substrate, 2.5 μM capture oligo, 2 mM ATP and 0-250 mM KCl. Control reactions containing no ATP were included in the assay.

Inhibition assays were performed by combining 4x nsp13/substrate/capture oligo (6 μL) stock solution in Assay Buffer with 2-fold ADP:AlF_3_ (prepared as a 2x 1:2 molar ratio stock) serial dilutions (12 μL) in helicase buffer in reaction wells. Reactions were initiated by the addition of a 4x ATP solution in helicase buffer. Final concentrations were 10 nM nsp13, 0.5 μM DNA/RNA substrate, 2.5 μM capture oligo, 2 mM ATP and 0-500 μM ADP:AlF_3_. Control reactions containing no ATP and no ADP:AlF_3_ were included in the assays. Data were fit using a sigmoidal dose dependent equation:

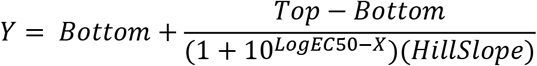

using Prism v.6.0 (GraphPad Software, Inc).

### Gel-based Helicase Activity Assays

Time-course experiments were performed by mixing recombinant nsp13 (20 nM) with ‘ partial duplex DNA or RNA substrates (100 nM) in the presence of ATP (2 mM) and capture oligo (1000 nM) in 80 μL reaction volumes. Reactions were performed in clear 200 μL tubes at 30°C in a temperature controlled aluminum block. Nsp13/substrate/capture oligos were prepared as a combined 4x stock solution in helicase buffer (20 mM HEPES, pH 7.5, 25 mM KCl, 5 mM MgCl_2_, 1 mM TCEP, 0.005% triton X100 and 0.1 mg/mL BSA), and 20 μL was added to 40 μL helicase buffer in the reaction tube. Reactions were initiated by the addition of 20 μL of a 4x ATP stock solution in helicase buffer. 10 μL reaction time-points were taken at 1, 2.5, 5, 10, 20, 30 and 60 min, and quenched by the addition of 10 μL of a 100 mM EDTA, 20% glycerol solution. Time-point samples (5 μL) were resolved by 10% native-PAGE at room temperature, 200 V for 25 min. The gels were scanned using a Typhoon Trio Variable Mode Imager (GE Healthcare) in TAMRA dye mode to visualize fluorescence.

Variable KCl concentration assays were performed by mixing recombinant nsp13 (200 nM) with partial duplex DNA or RNA substrates (20 nM) in the presence of ATP (2 mM) and capture oligo (2000 nM) in 10 μL reaction volumes. Nsp13/substrate/capture oligo was prepared as a 4x stock solution in helicase buffer and 2.5 μL was added to 5 μL of 2x stocks of KCl in helicase buffer (2-fold serial dilutions). Reactions were initiated by the addition of 2.5 μL of a 4x ATP stock solution in helicase buffer and incubated at 37 °C for 30 min. Reactions were quenched and gels were performed as described above.

### Preparation of Substrate for Optical Tweezers Assay

The substrate consists of two 1.5-kbp-long biotinylated dsDNA handles annealed to an RNA hairpin containing a 180-bp stem and a tetraloop. A 20-nt single-stranded region at the 5’ flank of the hairpin serves as the nsp13 loading site (with a sequence of 5’- CAGAAAGAAAAACCGGAUCC-3’). The hairpin sequence was made by in vitro transcription of a DNA template used previously^40^ using the MEGAscript T7 Transcription Kit (Thermo Fisher). RNA was then purified using the MEGAclear Kit (Thermo Fisher). The DNA handles were generated by PCR using modified primers to generate the overhang for annealing to the hairpin-containing RNA. The forward primer for the 5’ overhang handle contains an inverted base that terminates Phusion DNA polymerase (with a sequence of 5’- CAACCATGAGCACGAATCCTAAACCT/ilnvdT/GCATAACCCCTTGGGGCCTCTAAACG-3’). The forward primer for the 3’ overhang handle contains inverted bases at its 5’ end that terminate Taq polymerase (with a sequence of 5’- /5InvdG//iInvdC//iInvdA//iInvdA//iInvdA//iInvdT//iInvdC//iInvdT//iInvdC//iInvdC//iInvdG//iInvdG//iIn vdG//iInvdG//iInvdT//iInvdT//iInvdC//iInvdC//iInvdC//iInvdC//iInvdA//iInvdA//iInvdT//iInvdA//iInvdC//iInvdG/TAGTCTAGAGAATTCATTGCGTTCTGTACA/3ddC/-3’). The reverse primers for the 5’ and 3’ overhang handles contain a biotin at their respective 5’ ends for bead attachment.

The annealed complex containing the 5’ overhang handle and the RNA hairpin was assembled by mixing 1 *μ*M 5’ overhang handles, 400 nM RNA hairpin in a total volume 20 *μ*L of TL buffer, incubated at 65 °C for 90s and cooled to 4 °C. TL buffer consisted of 50 mM HEPES pH 7.5, 55 mM KOAc, 6 mM Mg(OAc)2, and 1 mM DTT. A 10 *μ*l reaction containing 3.5 *μ*L of annealed complex and 1.6 *μ*L of 0.5% streptavidin coated polystyrene beads (2 *μ*m diameter, Spherotech) was incubated on ice for 30 min. Then the mixture was diluted with 1 mL TL buffer. A separate reaction containing 50 nM 3’ overhang handles and 1.6 *μ*L of beads was incubated and diluted as above.

To form a tether, a bead conjugated to5’ overhang handle complexes was immobilized on a pipette via suction and brought into close proximity to a second bead conjugated to 3’ overhang handles held in an optical trap. Upon hybridization of the 3’ overhang handle with the 5’ overhang handle/RNA hairpin complex, a tether was formed.

### Optical tweezers measurements

All single-molecule measurements were made using an optical tweezers instrument (“MiniTweezers”)^41^. The sample buffer consisted of 20 mM HEPES pH 7.5, 20 mM KCl, 5 mM MgSO4, 1 mM TCEP, 0.1 mg/mL BSA, and 0.005% triton X100. At the start of each experiment, a tether was formed in the sample chamber and its force-extension behavior was obtained by moving the optically trapped bead at a speed of 70 nm/s away from the second bead held in the pipette. Once a single tether was confirmed by the shape of the force-extension curve, nsp13 (0.1 nM) and ATP (1 mM unless otherwise stated) were injected into the chamber, and unwinding data were collected in a force-feedback mode to maintain a constant tension in the tethered molecule. Data were collected at 200 Hz. All experiments were conducted at 23 ± 1 °C.

Single-molecule data were analyzed in MATLAB (Natick, MA). The raw readout in nm was converted to base pairs by using the extensible worm-like chain model with parameters ^42^: persistence length 1 nm, contour length 0.59 nm per RNA nucleotide, and stretch modulus 1000 pN. Velocity was determined by linear fitting to the time-distance data (start to end of a processive event). Processivity was determined as the maximum distance traveled prior to detachment. Event frequency was determined by counting the number of independent unwinding events divided by the total recording time for a given tether.

## SUPPLEMENTARY INFORMATION

Four supplemental figures

## ACKNOWLEDGEMENTS

We thank members of the Kapoor Lab as well as the Seth Darst and Elizabeth Campbell Labs for useful discussions. T.M.K is grateful to the NIH/NIGMS for funding (R35 GM130234-01). S.L. is supported by the Robertson Foundation, Sinsheimer Foundation, and NIH (DP2HG010510). K.J.M. is supported by a National Cancer Institute K00 Fellowship (K00CA223018). We are also grateful to the Pels Center for Biochemistry and Structural Biology for support.

## AUTHOR CONTRIBUTIONS

K.J.M. and X.C. performed single-molecule assays and processed data. P.M.M.S. performed bulk helicase assays. M.G. expressed and purified proteins with aid from K.J.M. and S.E.W. M.G. and P.M.M.S. performed DSF assays. K.J.M. performed fluorescence anisotropy assays. A.A. provided key reagents. S.L. and T.M.K. supervised research. K.J.M., P.M.M.S., M.G., S.L., and T.M.K. wrote the manuscript with input from all authors.

