## Supplementary material for "Force-dependent stimulation of RNA unwinding by SARS-CoV-2 nsp13 helicase": Supp Material

**This PDF file includes:**

**Figs. S1 to S4**

**Supplementary Figures**

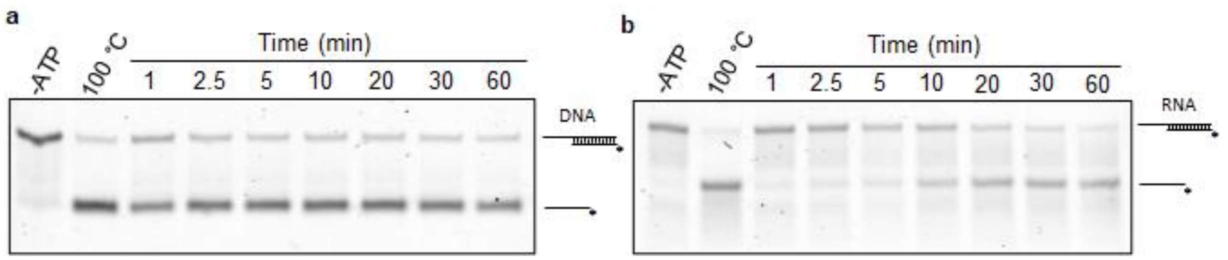


**Figure S1. Gel-based helicase assay time courses.**

a. Helicase assay time-course shows that SARS-CoV-2 nsp13 can unwind partial duplex DNA with a 5' overhang. TAMRA label is marked with an asterisk. Heat denatured and no ATP controls are indicated. Fluorescence was visualized using a Typhoon Trio Variable Mode Imager in TAMRA dye mode.

b. Helicase assay time-course shows that SARS-CoV-2 nsp13 can unwind partial duplex RNA with a 5' overhang. TAMRA label is marked with an asterisk. Heat denatured and no ATP controls are indicated. Fluorescence was visualized using a Typhoon Trio Variable Mode Imager in TAMRA dye mode.

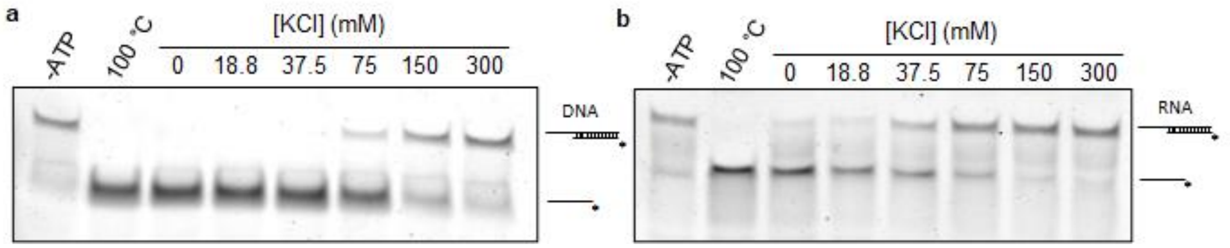


**Figure S2. Salt-dependence of nsp13 helicase activity in gel-based assays.**

a. Helicase assay shows the effect of salt concentration on SARS-CoV-2 nsp13 DNA helicase activity. TAMRA label is marked with an asterisk. Heat denatured and no ATP controls are indicated. Fluorescence was visualized using a Typhoon Trio Variable Mode Imager in TAMRA dye mode.

b. Helicase assay shows the effect of salt concentration on SARS-CoV-2 nsp13 RNA helicase activity. TAMRA label is marked with an asterisk. Heat denatured and no ATP controls are indicated. Fluorescence was visualized using a Typhoon Trio Variable Mode Imager in TAMRA dye mode.

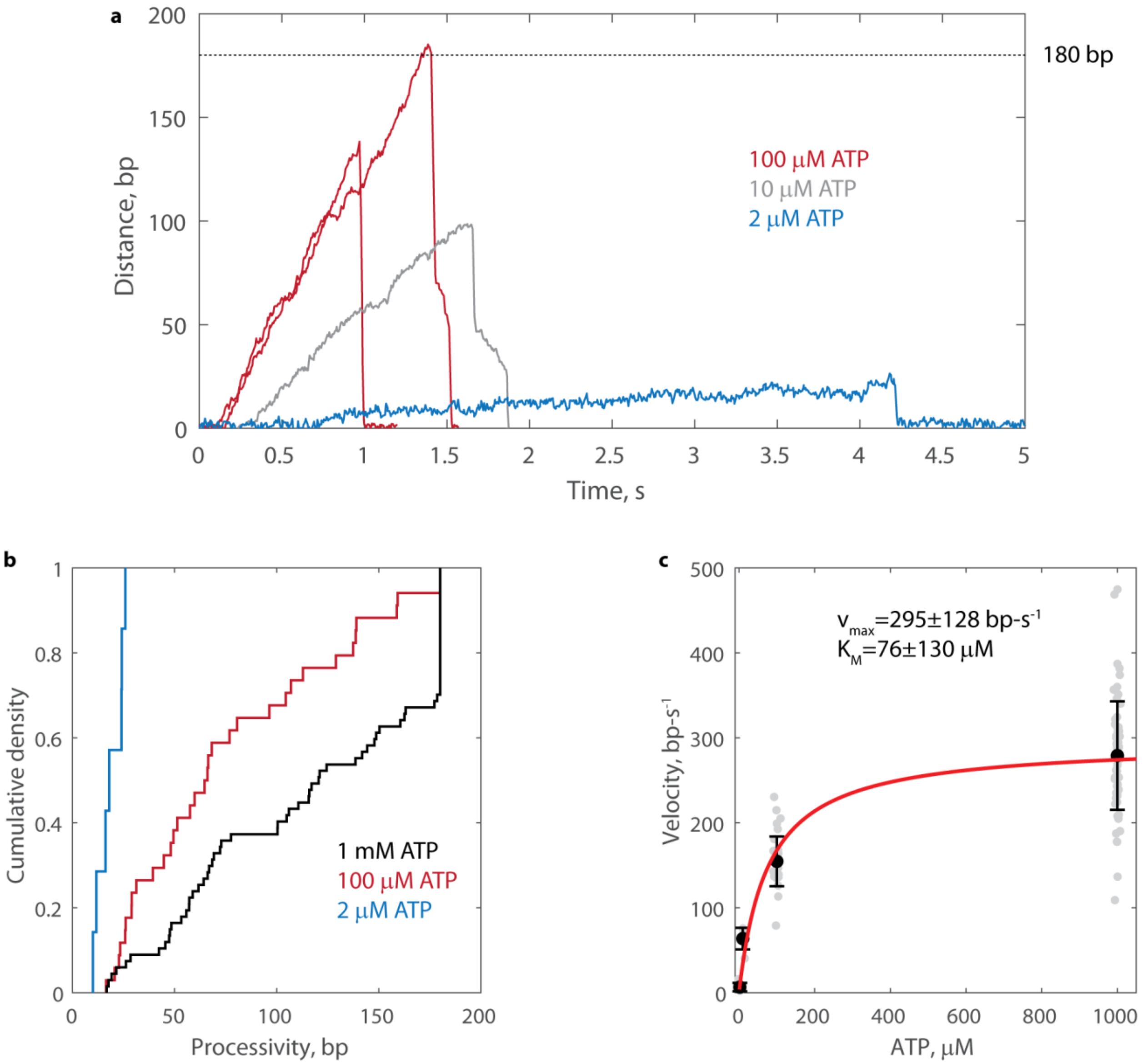


**Figure S3. Single-molecule measurements of nsp13 at 18 pN and various ATP concentrations**

a. Example traces of nsp13 unwinding activity on the same RNA hairpin structure as Fig. 3, but with different amounts of ATP.

b. The processivity distributions for nsp13 at 2 μM ATP (n=7), 100 μM ATP (n=34), and 1000 μM ATP (n=67).

c. The velocities of nsp at 2 μM ATP (n=7), 10 μM ATP (n=6), 100 μM ATP (n=34), and 1000 μM ATP (n=67). Gray dots show individual measurements, black dots show mean ± standard deviation. Red line shows fit to the Michaelis-Menton equation $v(x) = \frac{v_{max}x}{K_M+x}$, with fitted parameters (fit ± 95% confidence intervals) shown inset.

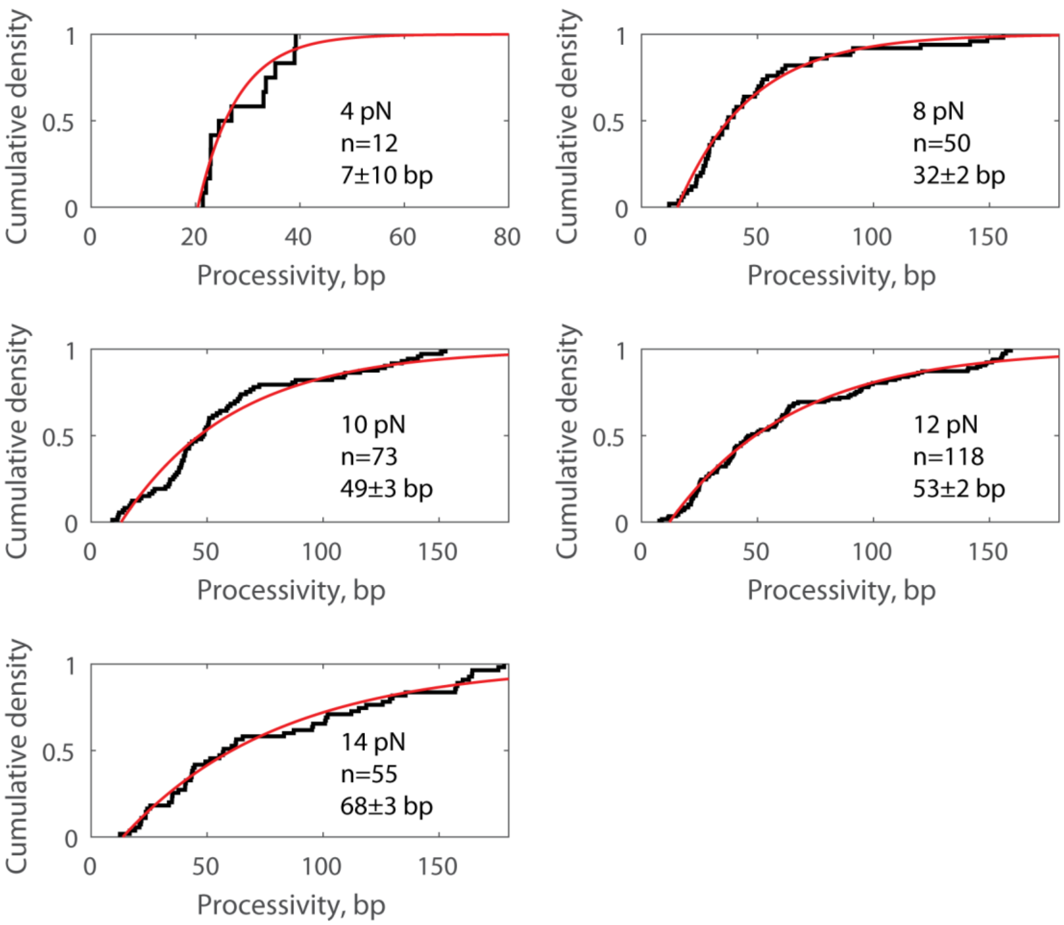


**Figure S4. Single-molecule processivity data fitted to single exponentials**
Each panel shows the processivity distribution of nsp13 at 1 mM ATP and the indicated force, replotted from Fig. 4c. Red line are fits to an offset exponential $F(x) = 1 - exp(\frac{-(x-x_{min})}{p})$ where $x_{min}$ denotes the smallest possible processivity measurement and p denotes the average processivity. Inset values show fitted values for p are shown as fit ± 95% confidence intervals.
